# Genomic in vitro transcription and Nanopore direct RNA sequencing of a human B-Lymphocyte cell line

**DOI:** 10.1101/2025.04.25.650674

**Authors:** Talia Tzadikario, Stuart Akeson, Sally Dorer, Neda Ghohabi Esfahani, Andrew J Stein, Pooria Daneshvar Kakhaki, Urja Choudhary, Kshitij Amar, Miten Jain

**Affiliations:** Department of Bioengineering, Northeastern University, Boston, MA, USA; Department of Physics, Northeastern University, Boston, MA, USA; Khoury College of Computer Sciences, Northeastern University, Boston, MA, USA

**Keywords:** Direct RNA Sequencing, Modification calling false positive, Genomic in vitro transcription, RNA Modification Negative Control Benchmarking

## Abstract

Genomic DNA used as a template for in vitro transcription of RNA can serve as a true negative control for benchmarking RNA modification detection by Nanopore direct RNA sequencing (DRS) models. We generated DRS data for in vitro transcribed (IVT) RNA composed of canonical nucleotides using genomic DNA from a human cell line. We applied Dorado modification calling models to these data, and calculated 9-mer specific false-positive rates for eight RNA modifications as a comparison point for future development of RNA modification models.

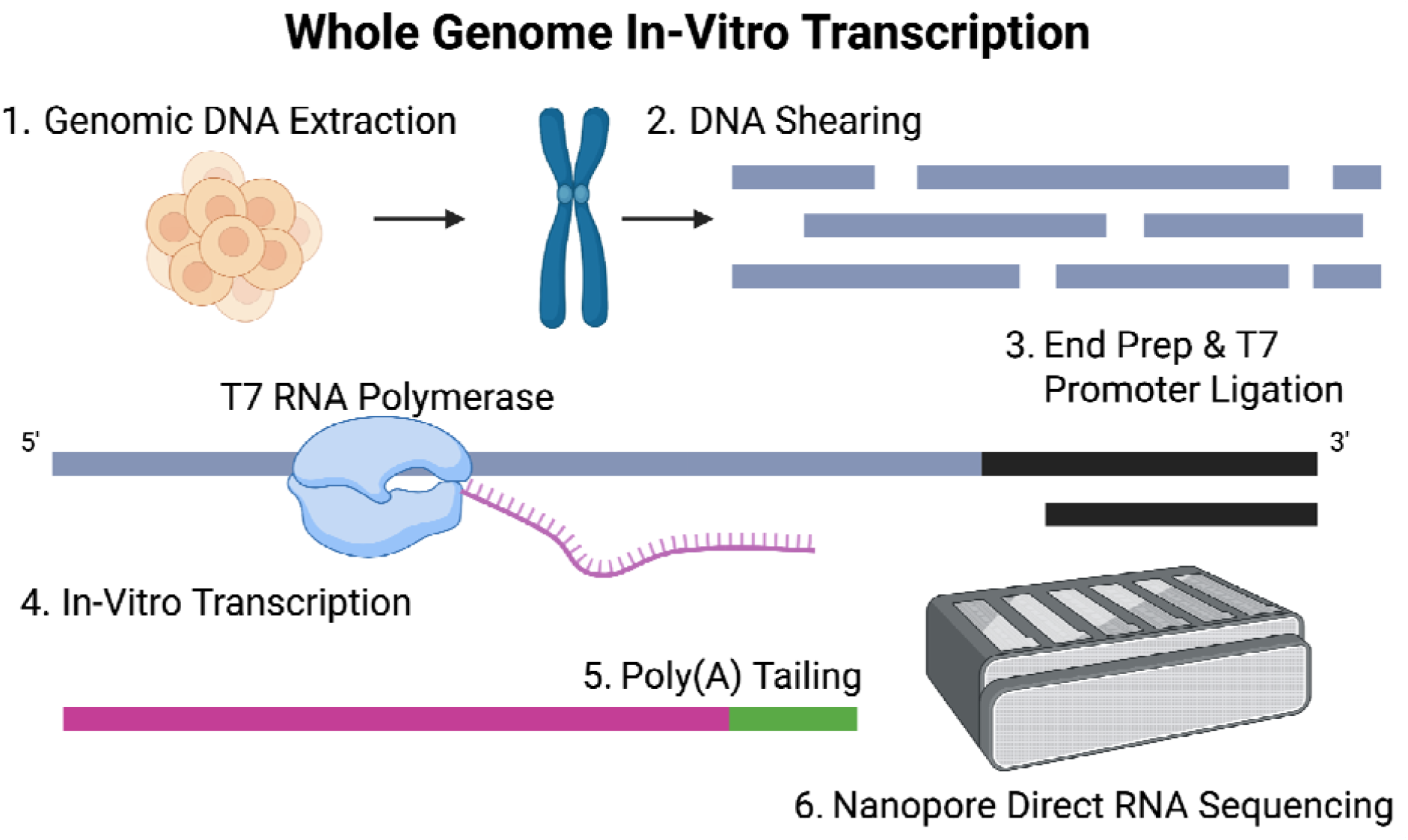

## Context

RNA modifications are chemical alterations to RNA nucleotides that have been shown to impact RNA function, structure, and half-life^1^. Over 170 RNA modifications have been identified to date across multiple RNA classes^2–5^. Research based on these observations has uncovered a host of biological implications from the modification of RNA molecules^5–7^. Additionally, alterations in the presence or absence of RNA modifications have been linked to disease pathology^1,8^. Detecting RNA modifications in a site and sequence-specific manner has been a challenging area for the field.

Nanopore direct RNA sequencing (DRS) offers a unique method of studying RNA modifications. DRS measures the disturbance of ionic current through a nanopore as an RNA strand translocates, providing a direct measurement of the native molecule, including RNA modifications present on the strand^9,10^. In 2024, Oxford Nanopore Technologies (ONT) introduced a new RNA chemistry with higher throughput and, when coupled with a new basecalling algorithm, improved accuracy (98%+)^11^. Additionally, this new software release for Dorado basecalling included ionic current-based RNA modification detection algorithms for all sequence contexts. Currently, Dorado includes eight RNA modification identification models: N6-methyladenosine (m^6^A), Inosine, N2-methyladenosine (2’-OMe-A), 5-methylcytosine (m^5^C), N2-methylcytosine (2’-OMe-C), N2-methylguanidine (2’-OMe-G), N2-methyluridine (2’-OMe-U) and Pseudouridine (Ψ)^11,12^. It is imperative that new RNA modification calling models are thoroughly benchmarked before their results are used to inform biological conclusions.

In this data note, we sequence all-canonical RNA generated by transcribing genomic DNA and analyze its use as an all context true-negative IVT dataset of all possible sequence contexts of length 9 (9-mers). Using this true negative data set we show that the ONT Dorado RNA modification models have 9-mer sequence specific false-positive rates ranging from less than 0.1% to 21.74%. This dataset provides a useful and easily accessible negative-control for benchmarking both future iterations of Dorado basecalling models as well as new RNA modification calling models developed by the community.

## Methods

### DNA extraction

Genomic DNA was extracted from a frozen pellet of 5 million GM12878 lymphoblastoid cells using the Qiagen Puregene Kit (Qiagen, Hilden, Germany), following the manufacturer’s protocol for cultured cells^13^. Briefly, cells were lysed in cell lysis buffer with Proteinase K at 37□°C until complete dissolution of cell material. Following RNase A treatment, proteins were precipitated using the provided precipitation buffer. DNA was precipitated with 100% isopropanol, and the resulting DNA pellet was resuspended in Tris-EDTA (TE) buffer and stored at 4□°C.

### Whole genome shearing, adapter ligation, in-Vitro Transcription and Poly(A) tailing

We took 4µg of the extracted DNA and sheared it to 6kb length using the Covaris g-Tube (Covaris 520079) by loading it and centrifuging it at 13.2k RPM for 1 min for each side three times using an Eppendorf 5424 centrifuge with rotor FA-45-24-11. Following this, we dA-tailed and repaired the DNA using NEBNext® Ultra™ II End Repair/dA-Tailing Module (E7180S) and NEBNext® FFPE DNA Repair Mix (E7180S) at room temperature for 5 minutes and 65°C for 5 minutes. This was followed by a 1X SPRISelect clean (B23317).

We proceeded with ligating our T7 Adapter to the sheared DNA. Briefly, this T7 Adapter was prepared by hybridizing in TNE (10mM Tris-HCl pH 8.0, 100mM NaCl, 1mM EDTA), 10µM 5’ - CCC TAT AGT GAG TCG TAT TAA TTT CGA - 3’ and 10µM 5’ - T GGG ATA TCA CTC AGC

ATA ATT AAA GCT - 3’, and heated to 75°C for 2 minutes, reduced to 25°C at a rate of 2°C per minute, and then rapidly cooled to 4°C before storage or use. The sheared and prepared DNA was ligated to the T7 Adapter at room temperature over 15 minutes in the presence of 2,000U of T4 DNA Ligase (NEB E7180S) and 10µM T7 Adapter. This was followed by a 2X SPRISelect clean (B23317).

1µg of the ligated DNA was used with the HiScribe® T7 Quick High Yield RNA Synthesis Kit (NEB E2050S) following the indicated protocol, except that transcription happened overnight. After, the resulting RNA and DNA mixture was purified using the optional DNAse I treatment recommended by and supplied by the HiScribe® T7 Quick High Yield RNA Synthesis Kit (NEB E2050S). This was followed by a 2X SPRISelect clean (B23317). 10µg of this was polyadenylated (M0276S) for 1 minute at 37°C using the indicated protocol followed by another 2X SPRISelect clean (B23317).

### Library preparation and sequencing

1µg of the in vitro transcribed and Poly(A) tailed RNA was prepared for sequencing following the ONT SQK-RNA004 protocol and sequenced on the PromethION platform.

### Basecalling and modification calling

Basecalling was performed using Dorado 1.0.0 super accuracy mode and the rna004_130bps_sup@v5.2.0 @v1 modification models were used for all eight modifications. The following Dorado basecalling parameters were used to produce the unaligned bam with modification calls:

~~~
dorado basecaller sup --emit-moves --modified-bases pseU_2OmeU m5C_2OmeC inosine_m6A_2OmeA 2OmeG --modified-bases-threshold 0
~~~

### Identity and Coverage Analysis

Reads were aligned to the GRCh38 reference genome using minimap2:

~~~
minimap2 -ax map-ont --MD -y | samtools view -F 2308 | samtools sort
~~~

Read alignment length was calculated as the total span covered on the reference genome and positional identity was calculated as the proportion of matches to the sum of matches, mismatches, insertions, and deletions. Coverage of gencode gene loci and exons was calculated using a whole genome IVT coverage summary where every position with a minimum coverage of 3 was counted and bedtools:

~~~
bedtools coverage -a gencode.v49.annotation.gff3 -b coverage_bed_mincov3.bed
~~~

### False Positive Rate Identification

Per-read per-site base modification status was determined using the Modkit^14^ extract full command. Only positions where the target base matched the reference were considered. A false positive is when the modification confidence exceeds the selected modification threshold. The modification threshold for the figures was set to 0.7, the recommended minimum for the cutoff threshold. Each position was collated into the parent 9-mer sequence and a modified and canonical total was calculated based on the confidence for modification thresholds ranging from 0.7 to 0.99 by increments of 0.01. These results were tabulated and are available in the supplementary data or on github (**Supplementary Table 1)**.

## Data Description

### Data Validation and Quality Control

Three PromethION genomic IVT DRS experiments produced 8,883,294 aligned RNA strand reads with a median alignment identity of 98.67% and an aligned N50 of 690 base **(Supplementary Figure 1A)**. This resulted in a median coverage of 10 reads per base across the entire GRCh38 reference genome with 0.73% of non-”N” positions having a coverage of 0. This includes at least some coverage of 99.59% of gene bodies in gencode v49 with a minimum read depth of 3 and complete coverage of 93.55% of exons in gencode with a minimum read depth of 3. The median proportion of mismatches, insertions and deletions to aligned length per read were 0.0034, 0.0028, and 0.0048, respectively **(Supplementary Figure 1B)**. The global false-positive rate, calculated as the proportion of modification calls out of the total considered positions using the precalculated threshold, ranged from 0.001557% for 2’-OMe-A to 0.008565% for 2’-OMe-G **(Supplementary Table 1)**. Nanopore sequencing is susceptible to sequence-specific biases in the signal^12,15^. We selected a sequence window size of 9 (9-mer) to capture as much context as possible while ensuring sufficient depth to estimate false-positives rates as low as 0.05% for each 9-mer. The genomic IVT’s complete representation of all 9-mers allows for a robust analysis of the 9-mer specific false positive rate.

### False Positive Rates in Commonly Studied Enzyme Motifs

Modification studies often focus on enzyme-specific sequence contexts to help elucidate the role of the enzyme and allow for knockdown and knockout cell conditions. We began our 9-mer false-positive calculations by analyzing false-positive rates for biologically relevant modification enzyme consensus 5-mer motifs. Each 5-mer motif (**Figure 1A**). With the exception of m^5^C all of the motifs we analyzed were below a 2.5% false-positive rate. For m^6^A and Ψ motifs, all of the 9-mers fell below 2.5% false-positive rates. m^5^C showed a higher biologically relevant false-positive rate, scoring as high as 0.072 for the m^5^C motif GGCUCCAGG. Notably, the pseudouridine motifs of TRUB1 (GUUCN) and PUS7 (UGUAR) show false-positive rates well below the highest pseudouridine false-positive rate of 20.27%. While a relatively low false-positive rate for these motifs is promising, negative controls and orthogonal validation are imperative for drawing biological conclusions. Because of the “black box” nature of the Dorado RNA modification calling models, it is not possible to attribute specific signal attributes to the increased false-positive rate. This is especially true since the training data used for the models are not made publicly available.

**Figure 1.**
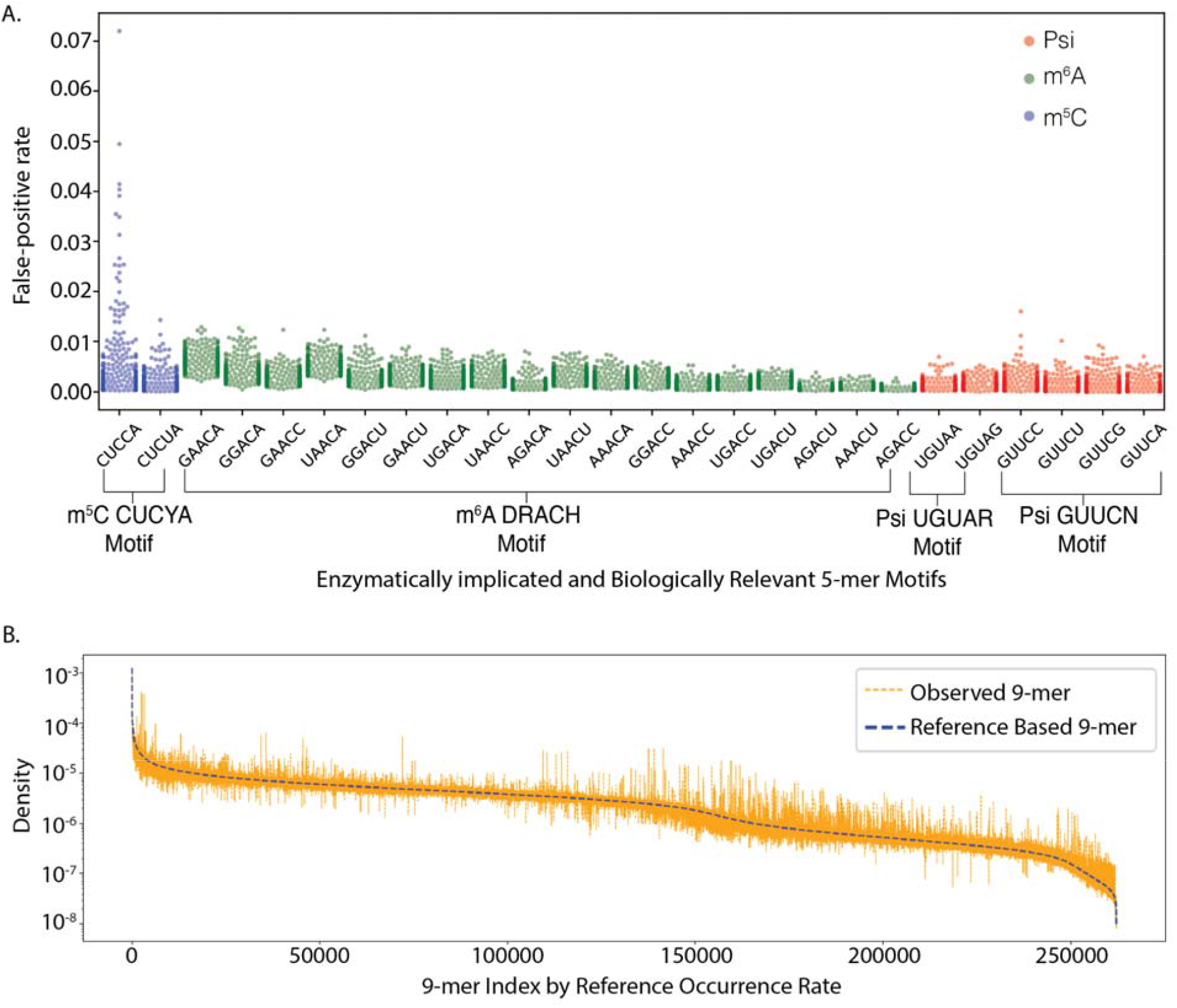
Performance and k-mer coverage analysis for a whole genome IVT derived RNA-based Nanopore dataset. **A**. False-positive rates for 5-mer motifs that have known RNA modifying enzymatic activity. **B**. Expected density of 9-mers based on the GRCh38 reference genome vs. the observed density of 9-mers in the whole Genome IVT dataset.

### All Context 9-mer Coverage

Dorado is not limited to biologically relevant motifs and claims all-context modification calling. We compared the frequency of observed 9-mers in the sequencing data to the expected distribution of 9-mers in the GRCh38 reference genome with the expectation that our observed 9-mer population should be similar to the reference genome (**Figure 1B**). The observed density approximates the expected distribution of 9-mers. The 9-mer with the lowest coverage was CGAUACGCG (218 reads), providing sufficient depth for estimating the false-positive rate of 9-mers. We note that coverage of all 9-mers can also be obtained from IVT generated from paired poly(A) RNA.

### 9-mer specific false-positive rates

For each modification, we calculated a per-9-mer proportion of false positive modification calls using a 0.7 confidence threshold **(Figure 2A-H)**. The 0.7 threshold was selected as a minimum credible threshold using the Modkit toolkit^14^, below this threshold, users are given a warning by Modkit and instructed to set a fixed threshold of 0.7. Inosine, 2’-OMe-C and 2’-OMe-A showed similar patterns of false positive distributions, with the top 5% of false-positive rates ranging from 1.40% to 7.53% for Inosine, 1.03% to 9.88% for 2’-OMe-C and 0.81% to 6.61% for 2’-OMe-A (**Table 1**). m^6^A, 2’-OMe-G, and 2’-OMe-U showed a heightened distribution of false-positive rates in the top 5 percentile with 9-mer specific maximums ranging from 10.19% to 14.69%, while m^5^C and Ψ showed more exaggerated pattern of false positive predictions, especially at the extrema, with the top 5% of false positives ranging from 3.57% to 21.74% and 1.41% to 20.27% respectively (**Table 1**). We additionally calculated the false-positive threshold for each modification in each appropriate 9-mer, ranging from a cutoff of 0.7-0.99 by increments of 0.01. This data is available on github^16^.

**Table 1.** Modification calling false-positive rates for subsets of up to the top 5^th^ percentile.

| False positive percentile | m<sup>6</sup>A | Psi | Inosine | m<sup>5</sup>C | 2'-OMe-G | 2'-OMe-C | 2'-OMe-U | 2'-OMe-A |
| --- | --- | --- | --- | --- | --- | --- | --- | --- |
| Chm 22 Single Position Max at minimum 10 coverage | 0.70 | 0.609 | 0.909 | 0.900 | 0.600 | 0.707 | 0.727 | 0.400 |
| 9-mer Max | 0.1019 | 0.2027 | 0.0753 | 0.2174 | 0.1469 | 0.0988 | 0.1447 | 0.0661 |
| 0.01 | 0.077 | 0.1385 | 0.0631 | 0.1814 | 0.1337 | 0.0836 | 0.1318 | 0.0494 |
| 0.1 | 0.0594 | 0.0875 | 0.0485 | 0.1225 | 0.1035 | 0.044 | 0.0853 | 0.0287 |
| 0.5 | 0.0433 | 0.047 | 0.0368 | 0.0838 | 0.072 | 0.0256 | 0.0562 | 0.0199 |
| 1 | 0.0357 | 0.0349 | 0.0301 | 0.0677 | 0.06 | 0.0206 | 0.047 | 0.0165 |
| 2 | 0.0295 | 0.0248 | 0.0232 | 0.0529 | 0.0488 | 0.0159 | 0.0371 | 0.0128 |
| 3 | 0.0257 | 0.0196 | 0.0187 | 0.0446 | 0.0422 | 0.0134 | 0.0314 | 0.0106 |
| 4 | 0.0233 | 0.0163 | 0.0159 | 0.0396 | 0.0371 | 0.0116 | 0.0269 | 0.0093 |
| 5 | 0.0215 | 0.0141 | 0.014 | 0.0357 | 0.0332 | 0.0103 | 0.0234 | 0.0081 |
| Portion of 9-mers with no false-positive | 0.0180 | 0.0165 | 0.0104 | 0.0013 | 0.0694 | 0.1563 | 0.0052 | 0.0089 |

**Figure 2.**
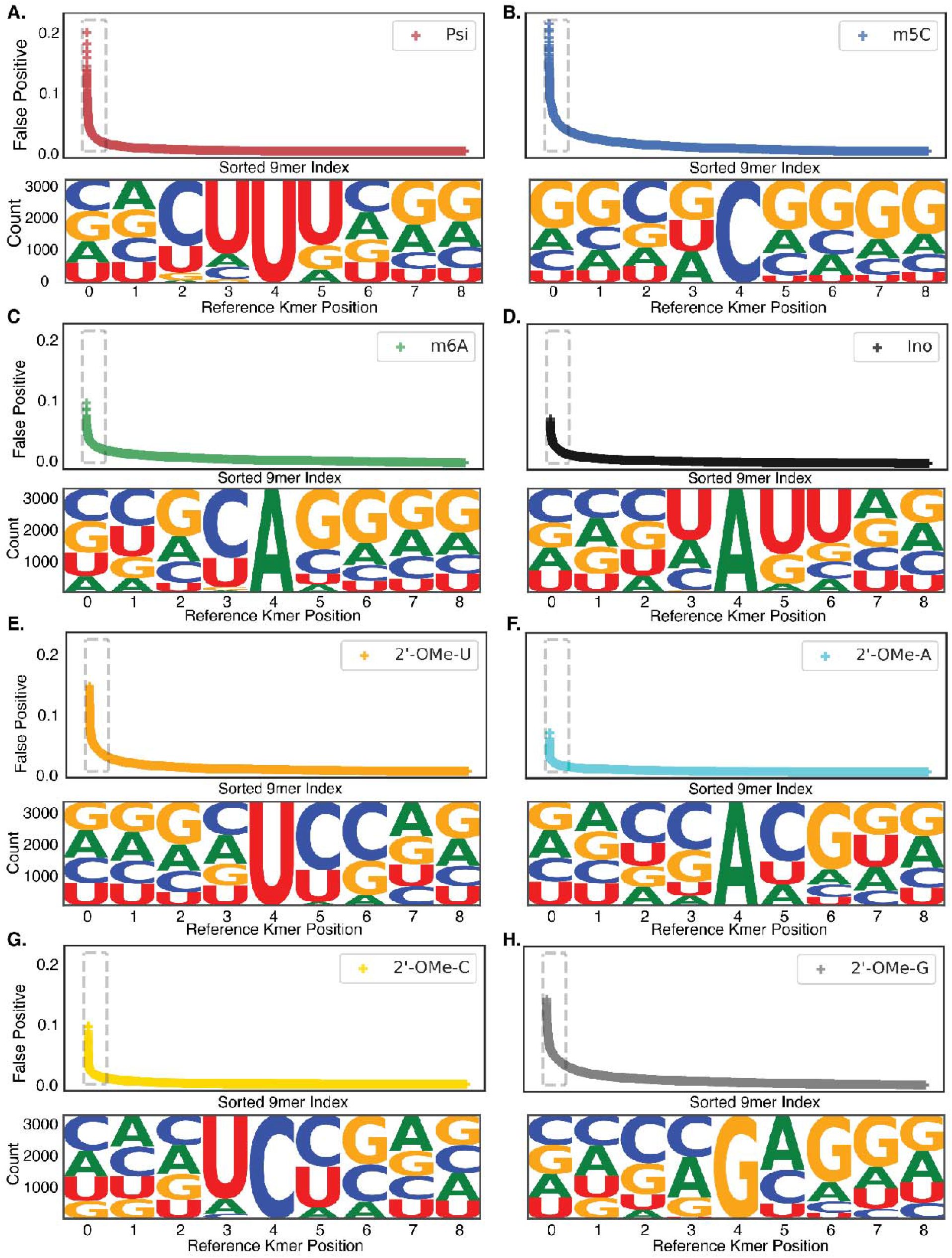
9-mer specific false-positive rate for 8 Dorado called modifications. **A.** False-positive rates per 9-mer for Ψ sorted from highest to lowest. The distribution of bases in the highest 5% of false positive 9-mers. **B**. False-positive rates per 9-mer for m^5^C sorted from highest to lowest. The distribution of bases in the highest 5% of false positive 9-mers. **C**. False-positive rates per 9-mer for m^6^A sorted from highest to lowest. The distribution of bases in the highest 5% of false positive 9-mers. **D**. False-positive rates per 9-mer for Inosine sorted from highest to lowest. The distribution of bases in the highest 5% of false positive 9-mers. **E**. False-positive rates per 9-mer for 2’-OMe-U sorted from highest to lowest. The distribution of bases in the highest 5% of false positive 9-mers. **F**. False-positive rates per 9-mer for 2’-OMe-A sorted from highest to lowest. The distribution of bases in the highest 5% of false positive 9-mers. **G**. False-positive rates per 9-mer for 2’-OMe-C sorted from highest to lowest. The distribution of bases in the highest 5% of false positive 9-mers. **H**. False-positive rates per 9-mer for 2’-OMe-G sorted from highest to lowest. The distribution of bases in the highest 5% of false positive 9-mers.

For all 8 modifications, we can observe a dominant motif or partial motif for the highest false-positive rates. The top 5% of false positive sites for 2’-OMe-G, 2’-OMe-A, 2’-OMe-U, m^5^C, m^6^A, Ψ, and inosine are all moderately enriched in repetitive sequences of a single nucleotide around the modified site. This is expected due to complications of resolving homopolymeric sequences using nanopores^17^ (**Figure 2A-H**). Contrary to this observation, 2’-OMe-C’s top 5% of false positives are dominated by UCU and UCA motifs at positions 4-6 in the 9-mer. Researchers using Dorado modification calling should analyze homopolymeric motifs with additional scrutiny. Context aware analysis can help to identify positions with potentially inflated rates of modification in biological samples. We compared 9-mers for those bases that have multiple modifications: A, U, and C. Across all three compared canonical bases, there was a very low level of overlap between high false positive 9-mers (**Supplementary Figure 2A-C**). This suggests that false-positive rates are both 9-mer and modification specific.

### 9-mer false-positive rate variability between batches

Additionally, we compared three replicates of whole-genome IVT sequencing to document whether some 9-mers experienced higher levels of false positive standard deviation across runs. The highest standard deviation of false-positive rate we observed was 0.0296 for the 9-mer CCGAGTCGT. The 95th percentile of standard deviation of false-positive rate in our dataset is 0.0027, while the 5th percentile is 4.6269×10^-5^. The skew of the distribution is 4.769 with a mean of 0.00075 and an inter-quartile range of 0.0007 (**Supplementary Figure 2D)**. The batch to batch variability in 9-mer false positive rates, while small, suggests that negative controls like paired IVT should be further augmented with additional orthogonal validation steps.

### Basecalling uncertainty and false positive modification calls

Another possible explanation for false-positive calls is a lack of certainty from the basecaller in the initial canonical base call. It should be specifically noted that the confidence of the basecaller is not the same as accuracy; a basecall can have a high confidence and be incorrect. We hypothesized that bases with lower confidence values might be more prone to false positive modification calls. This would materialize as a negative correlation between base confidence and the modification model’s confidence of calling a modification. When we plotted this relationship for reads mapping to chromosome 22, we observed that there was no indication of a negative correlation (**Supplementary Figure 2E**, Non-zero mod quality; Spearman-r = −0.094).

### Reuse Potential

In this data note, we provide a dataset of the ionic current of canonical, all-context 9-mers. Creating RNA from genomic DNA helps to represent low-coverage regions that researchers want to analyze, which could be difficult to otherwise achieve. This data is also available for ionic current model training for purposes beyond modification calling.

As Nanopore DRS RNA modification calling models are used in biological research, it is important that their limitations are communicated clearly. With the dataset presented here, we demonstrated that contemporary RNA modification models have sequence specific false-positive rates when grouped into 9 nucleotide windows. While this is not a new discovery in the field of RNA modification detection using nanopores, this dataset provides an all-context negative control for benchmarking future iterations of Dorado basecalling models as well as new RNA modification calling models.

## Data Availability

We have shared the links to the data as well as the analysis code, files and pre-computed false-positive rates are available at https://github.com/genometechlab/gIVT. The basecalled data are available via ENA (PRJEB102644) and signal data are available via AWS (https://s3.amazonaws.com/nanopore-human-wgs/index.html?prefix=rna/NU_RNA004/gIVT/).

## Declarations

M.J. is a paid consultant to ONT. M.J. has received reimbursement for travel, accommodation, and conference fees to speak at events organized by ONT. Other authors declare no competing interests.

## Acknowledgements

We would like to acknowledge Sofie Salama and Kristof Tigyi at UC Santa Cruz for providing us with GM12878 cells. Additionally we would like to acknowledge Biorendr for providing the tools to construct the graphical abstract, the template for this figure can be found at *https://BioRender.com/1ok8q0b*

