## Supplementary Figures for "Genomic in vitro transcription and Nanopore direct RNA sequencing of a human B-Lymphocyte cell line"

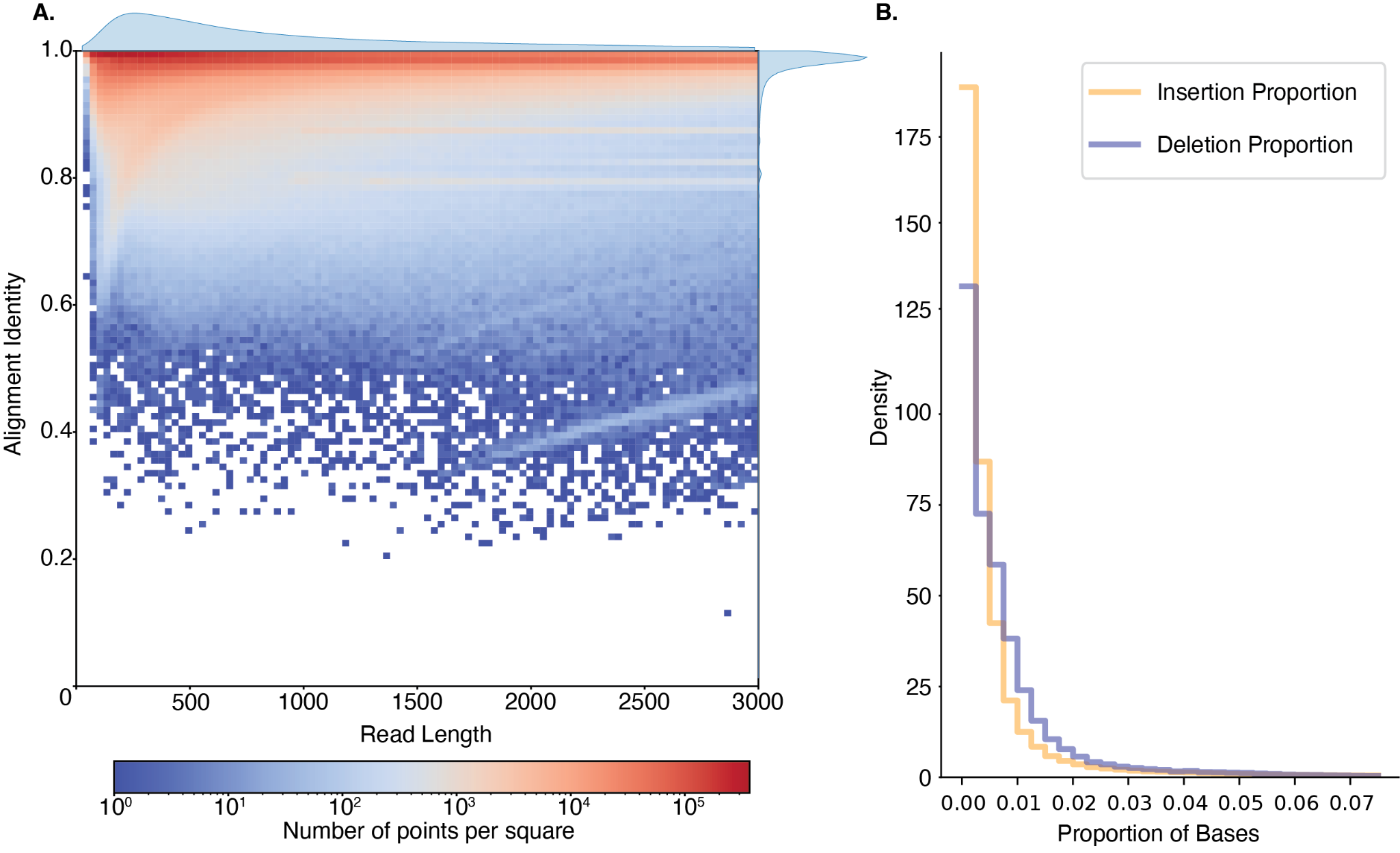

**Supplementary Figure 1. A.** Heatmap of alignment identity calculated as the proportion of bases matching the reference to the total number of matches, mismatches, insertions and deletions in the aligned region. **B.** Per-base proportion of insertions and deletions.

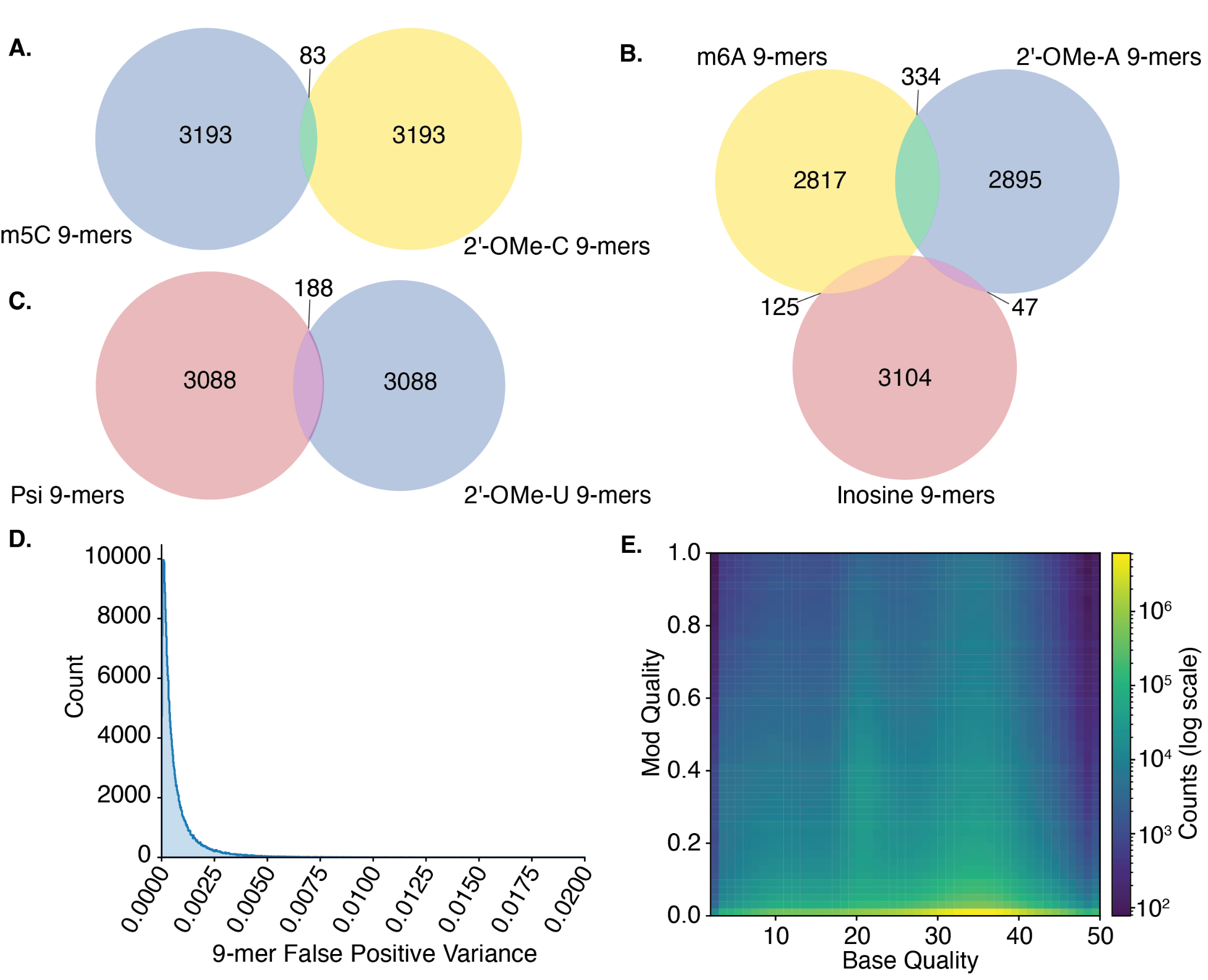

**Supplementary Figure 2.** **A.** Overlap of 9-mers in the top 5 percentile of false positives for m^5^C and 2’-OMe-C. **B.** Overlap of 9-mers in the top 5 percentile of false positives for m^6^C, Inosine and 2’-OMe-C. **C.** Overlap of 9-mers in the top 5 percentile of false positives for Pseudouridine and 2’-OMe-U. **D.** 9-mer specific false-positive variance between three technical replicates. **E.** Correlation between modification calls and base call confidence.

| **Supplementary Table 1**. Global false-positive rates for a minimum modification threshold of 0.7. | |
| --- | --- |
| Modification | Global false-positive rate |
| m^6^A | 0.005406 |
| Psi | 0.003592 |
| Inosine | 0.003442 |
| m^5^C | 0.006687 |
| 2'-OMe-G | 0.008565 |
| 2'-OMe-A | 0.001557 |
| 2'-OMe-C | 0.002961 |
| 2'-OMe-U | 0.005155 |
